# Single-cell analysis reveals an active and heterotrophic microbiome in the Guaymas Basin deep subsurface with significant heterotrophic inorganic carbon fixation

**DOI:** 10.1101/2023.11.06.565840

**Authors:** Nicolette R. Meyer, Yuki Morono, Anne E Dekas

## Abstract

The marine subsurface is a long-term sink of atmospheric carbon dioxide with significant implications for climate on geologic timescales. Subsurface microbial cells can either enhance or reduce the potential for the subsurface to sequester carbon, depending on their metabolic activity. However, the activity of subsurface microbes is rarely measured, leaving their role in biogeochemical cycling poorly characterized. Here, we used nanoscale secondary ion mass spectrometry to quantify anabolic activity in 3,203 individual cells from the thermally altered deep subsurface in the Guaymas Basin, Mexico (3–75 m below the seafloor, 0-14 C). We observed that a large majority of cells were active (83–100%), although rates of biomass generation were low, suggesting cellular maintenance rather than doubling. Mean single-cell activity decreased with increasing sediment depth and temperature, and was most strongly correlated with porewater sulfate concentrations. Intracommunity heterogeneity in cell-specific activity decreased with increasing sediment depth and age. Using a dual-isotope labelling approach we determined that all active cells analyzed at all depths were heterotrophic. We detected and quantified inorganic carbon assimilation by heterotrophs and found that it contributes on average at least 5% of total heterotrophic biomass carbon in this community. Our results therefore suggest that the deep marine biosphere at Guaymas Basin is largely active and contributes to subsurface carbon cycling primarily by assimilating organic carbon but also by mediating heterotrophic inorganic carbon fixation. Heterotrophic assimilation of inorganic carbon may be a small yet significant and widespread underappreciated source of labile carbon in the global subsurface.

**Importance:** The global subsurface is the largest reservoir of microbial life on the planet yet remains poorly characterized. The activity of life in this realm has implications for long-term elemental cycling, particularly of carbon, as well as how life survives in extreme environments. Here, we recovered cells from the deep subsurface of the Guaymas Basin and investigated the level and distribution of activity, the physicochemical drivers of activity, and the relative significance of organic versus inorganic carbon to subsurface biomass. Using a sensitive single-cell assay we find that the majority of cells are active, that activity is likely driven by availability of energy, and that while organic carbon supplies most cellular carbon, inorganic carbon also contributes. We additionally find that the inorganic carbon assimilation observed was mediated by heterotrophs, not autotrophs, highlighting the importance of this often overlooked mode of carbon assimilation in the subsurface and beyond.

## Introduction

Over 80% of the planet’s bacterial and archaeal biomass is estimated to be found within the global subsurface (Bar-On et al., 2018). With depth into the subsurface, life becomes progressively more challenging, reaching the energetic, thermal and pressure limits of life (Wagner and Kallmeyer 2014). These extreme conditions – particularly low energy availability – select for slower growing microorganisms that efficiently convert substrates into cellular energy (Morita 1997; Hoehler and Jørgensen 2013; Lever et al. 2015; Bradley et al. 2019) and may ultimately cause cells to resort to sporulation or minimal anabolic activity (Parkes et al. 2000; De Rezende et al. 2013; Bradley et al. 2020). We know that some cells in the marine subsurface are active, based on porewater geochemistry (D’Hondt et al. 2004), radiotracer experiments (Parkes et al. 1994, 2000, 2005; Schippers et al. 2005; Beulig et al. 2018, 2022; Heuer et al. 2020), stable isotope tracer experiments (Morono et al. 2011, 2020; Trembath-Reichert et al. 2017), and amino acid racemization models (Lomstein et al. 2012; Braun et al. 2017). Previous studies using single-cell approaches have broadly suggested 7 to 100% of deep subsurface cells are active (Morono et al. 2011, 2020; Trembath-Reichert et al. 2017). But intricacies of subsurface microbial activity remain poorly constrained, including how the proportion of active cells changes geographically and with physicochemical variables, how evenly activity is distributed within a community, and the role of active cells in carbon cycling. Constraining subsurface activity at the single-cell level can improve our understanding of the role of the deep biosphere in biogeochemical cycles as well as potentially life under extreme energy limitation more broadly.

Marine sediments are long-term sinks of atmospheric carbon dioxide with major climate implications on geological timescales. The relative rates of organic versus inorganic carbon fixation in subsurface cells has implications for the role of these communities in the biological pump, since the former counteracts carbon burial and the later can enhance it. Most archaeal and bacterial taxa that inhabit the deep biosphere are uncultured (Teske and Sørensen 2008; Steen et al. 2019), but metagenomic evidence provides insight to their metabolic potential. Previous work has demonstrated that genes for organic carbon degradation are widespread throughout the marine subsurface (Biddle et al. 2006; Lloyd et al. 2013; Orsi et al. 2013; Baker et al. 2015; Marshall et al. 2018; Orsi 2018; Zinke et al. 2019; Wasmund et al. 2021), suggesting widespread heterotrophy. However, hydrogenotrophic methanogens (Beulig et al. 2018) and some anaerobic methanotrophs (Kellermann et al. 2012) present in the subsurface are capable of assimilating inorganic carbon. In addition, heterotrophs can assimilate inorganic carbon through e.g. anaplerosis (Braun et al. 2021), but this is generally difficult to quantify in the environment and has never before been observed directly in the subsurface. Rate measurements are therefore needed to assess the relative importance of these processes.

The Guaymas Basin, Mexico, is an active rift system characterized by the emplacement of basaltic sills into organic rich sediments (Einsele et al. 1980; Von Damm et al. 1985). The thermal decomposition of organic matter produces reactive dissolved organic carbon (C_org_)(Lin et al. 2017), aliphatic and aromatic hydrocarbons (Simoneit and Lonsdale 1982; Kawka and Simoneit 1987; Didyk and Simoneit 1989), organic acids (Martens 1990), NH_4_^+^ (Von Damm et al. 1985), H_2_S and CO_2_ (Seewald et al. 1990). The mobilized products move upwards (Einsele et al. 1980; Lonsdale and Becker 1985), and can be a source of climate-warming CH_4_ and CO_2_ to the ocean-atmosphere system (Welhan 1988; Lizarralde et al. 2011). However, the products of thermal decomposition can also fuel deep subsurface microbial life, that use these substrates as metabolites, reducing the flow of greenhouse gases out of the marine subsurface (Teske et al. 2019a). Although activity has been widely studied in the surficial Guaymas Basin sediments (0– 30 cm) [e.g., (Elsgaard et al., 1994; Jorgensen et al., 1990; Kallmeyer & Boetius, 2004; Møller et al., 2018; Weber & Jørgensen, 2002)], microbial anabolic activity in the deep subsurface (>1 m) has not been explored at this site.

The steep temperature and geochemical gradients in the Guaymas Basin deep subsurface provide a particularly valuable environment to investigate microbial activity and its physicochemical controls. We set up a series of stable isotope incubations using sediment obtained during the International Ocean Discovery Program (IODP) expedition 385 via ocean drilling from 3–75 m below the seafloor (mbsf) at Site U1550, in the northern axial trough (Figure 1). We measured the incorporation of ^15^N-ammonium and ^13^C-bicarbonate over time using nano secondary ion mass spectrometry (nanoSIMS) to quantify cell-specific anabolic activity and investigate the single-cell heterogeneity of activity within microbial communities. Measures of microbial activity were interpreted in their physiochemical contexts – particularly geochemical conditions and temperature regimes – to understand the drivers of subsurface microbial activity. Additionally, leveraging a recently developed approach to identify autrotrophs and heterotrophs using cell-specific dual-stable isotope labelling, we characterize the dominant carbon assimilation strategy (heterotrophy versus autotrophy) for each cell, and determine the overall significance of inorganic carbon assimilation by heterotrophic cells.

**Figure 1:**
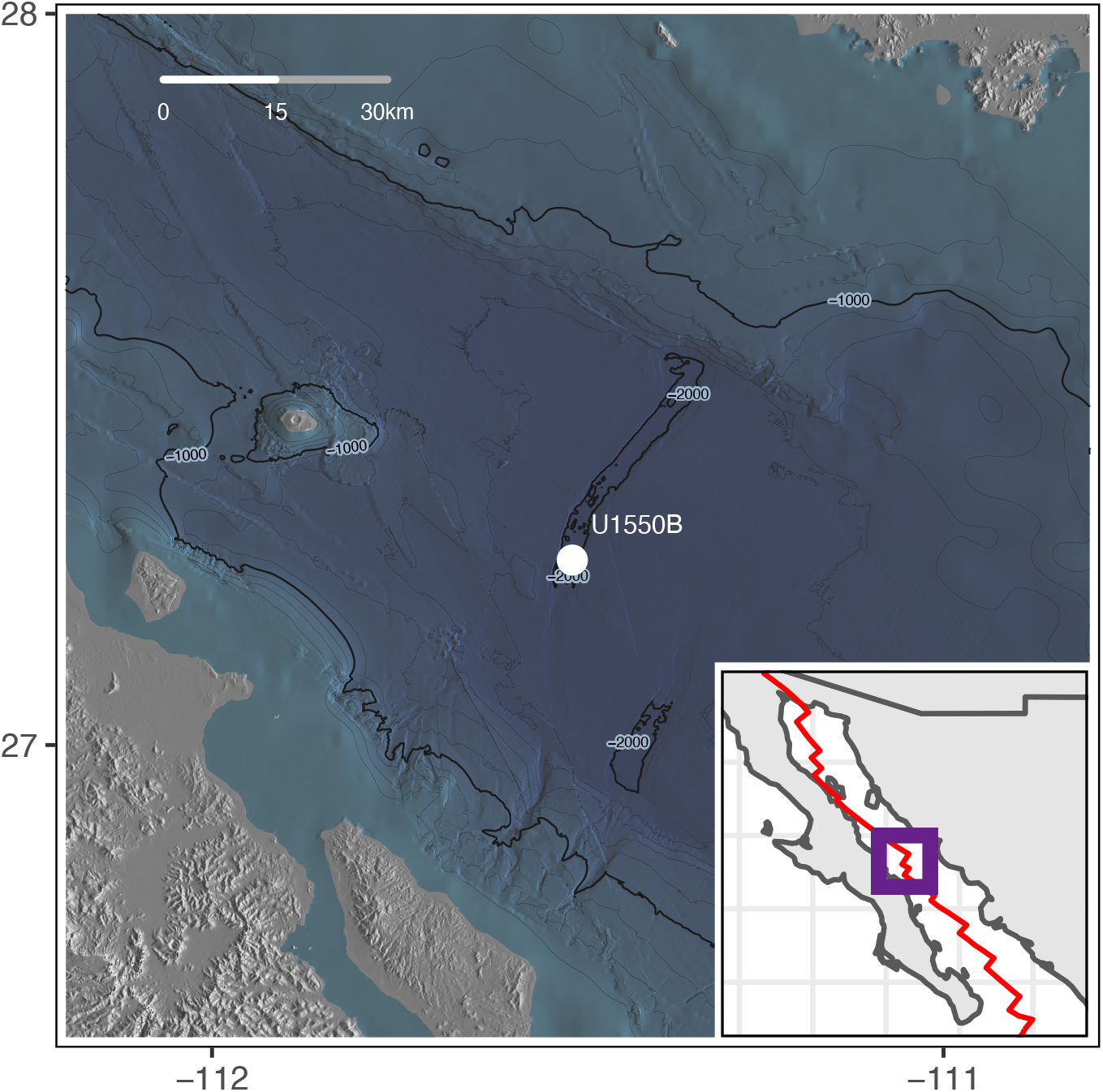
Map of Guaymas Basin showing location of U1550B (solid circle, 27° 15.1704’N, 111° 30.4451’W). Contour lines are 200 m apart. The inset shows the Gulf of California with the spreading centers and transform faults shown in red. Base map from (Ryan *et al*., 2009).

## Methods

### Sample collection

Marine sediment cores were collected on the D/V JOIDES Resolution on expedition 385 (Guaymas Basin Tectonics and Biosphere) from September 16^th^ 2019 to November 16^th^ 2019 in the Gulf of California, Mexico (Teske et al. 2021a) (Figure 1 and Table 1). Cores from Hole U1550B were collected using an advanced piston corer (APC), were immediately sectioned on the receiving platform, and capped to minimize oxygen exposure and contamination. The 30 cm long samples were kept in their capped core-liners, sealed in N_2_ gas-flushed mylar bags and stored at 4°C for 39–40 days until incubation setup. Porewater, solid geochemistry, and porosity was analyzed shipboard using established IODP protocols (http://www-odp.tamu.edu/publications/pubs_tn.htm).

**Table 1:**
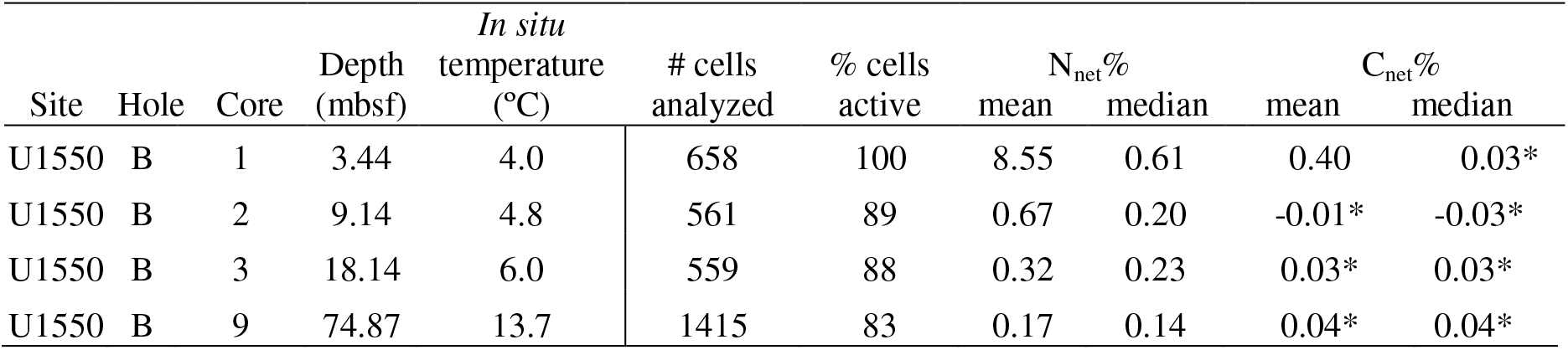
Sampling information and summary of nanoSIMS results. Astericks indicate values below the single-cell detection limit for significant isotope enrichment.

### Contamination control

The APC coring systems are dedicated to recovering samples with minimal contamination from drilling fluid (seawater). Perfluorocarbon tracers (PFTs) were deployed downhole for all cores to monitor drilling fluid contamination (Teske et al. 2021a) and revealed minimal contamination (across all holes and cores, 0.1 ppb in the external 1 cm, 0.0 ppb in the center, and 0.0 ppb halfway between the center and the outer rim; IODP LIMS database accessed at https://web.iodp.tamu.edu/). The outer ≥1 cm rind of sample was removed before microcosm setup.

### Microcosm experiments

Sediment (30 cm) from cores 1H-3 (3.4 mbsf), 2H-3 (9.1 mbsf), 3H-3 (18.1 mbsf), 9H-3 (74.9 mbsf), and 17F-3 (132 mbsf) were mixed with a modified, sterile anaerobic seawater medium (DSMZ medium 504) in a 1:2 ratio (sediment:media) in an anaerobic chamber. The medium contained 10% of the trace element mixture, selenite-tungstate solution, and vitamin mixture as specified in the recipe. 35 mL of sediment slurry was added to 50 mL serum vials, sealed with NaOH pre-treated black butyl rubber stoppers (Geo-microbial technologies, Ochelata, OK, USA) and aluminum crimp caps, and the headspace exchanged using argon. Incubations were amended with ^15^N-ammonium (final concentration: 3.34 mM, 99 at%; Cambridge Isotopes, NLM-467, Lot I-19633L), ^13^C-sodium bicarbonate (5.71 mM at%; Cambridge Isotopes, CLM-441, Lot PR-30836), and ^34^S-sulfate (2.0 mM; 80.4 at%; Sigma Aldrich, 718882, Lot MBBC1790). Incubations were over-pressured to 20 psi using argon and subsampled at ∼7, ∼14, ∼30, ∼90, and 304 days. The slurry was fixed in 0.2 µm filter-sterilized 2% paraformaldehyde (Electron Microscopy Sciences, Hatfield, PA, USA) in 1x phosphate buffered saline solution (PBS). Fixed samples were incubated overnight at 4°C and subsequently washed with PBS and ethanol, as previously described by (Dekas and Orphan 2011). After washing, the samples suspended in 100% ethanol were stored at −20°C until further processing.

### Cell extraction, counting, and sorting

Fixed cells from the 304 day-time point were extracted from the sediment slurries by density separation. A portion (10%) of the extracted material was trapped onto 0.2 μm polycarbonate filters and used for microscope cell counts for the determinate of cell density. The remainder (90%) was subjected to fluorescence-activated cell sorted (FACS) for further separation from the mineral matrix. Both the density separations and cell sorts were carried out in the clean-booth and clean-room facilities at the Kochi Institute for Core Sample Research, Japan Agency for Marine-Earth Science and Technology (JAMSTEC) as previously described by (Morono et al. 2013, 2020). The final point was selected for sorting and NanoSIMS analysis due to low microbial activity rates in the marine deep subsurface and to increase the chance of isotope label incorporation. Cells fixed immediately after collection were used to determine initial cell density. Cells were sorted onto custom ITO-coated 0.2 μm polycarbonate filters (Isopore GTBP02500; Millipore) (Morono et al. 2020). The filters were stored at −20°C until nanoSIMS analysis.

### nanoSIMS

Individual cells on ITO coated filters were analyzed using a CAMECA NanoSIMS 50 L (Stanford Nano Shared Facility, Stanford, CA, USA). A Cs^+^ primary ion beam (4 pA) with a nominal spot size of 100–200 nm was used to raster over cells of interest. Seven distinct secondary ion species were detected (^12^C_2_ ^-^, ^13^C^12^C^-^, ^14^N^12^C^-^, ^15^N^12^C^-^, ^31^P^-^, ^32^S^-^, and ^34^S^-^ using electron multipliers. The instrument was operated with a mass resolving power of at least 6500 to resolve isobaric interferences between ^13^C^12^C^-^ and ^12^C^1^H^-^, ^13^C_2_^-^ and ^12^C^14^N^-^, as well as ^11^B^16^O^-^ and ^12^C^15^N^-^ (Pett-Ridge and Weber 2022). At least thirty raster images were acquired at 256 × 256 pixel resolution for >30 min with a dwell time of ∼1000 μs/pixel. The base analysis chamber pressure was maintained at 10^-9^ torr. Non-isotope enriched *Methanosarcina acetivorans* cells with known isotope ratio mass spectrometry (IRMS) isotope composition were used as bracketing standards and to correct isotope ratios for instrumental mass fractionation (Meyer et al. 2017; Pett-Ridge and Weber 2022). Image processing was performed using Look@NanoSIMS software (Polerecky et al. 2012). Images were accumulated and aligned and regions of interest (ROIs) were drawn using the ^32^S^-^ and ^14^N^12^C^-^ images as bases. ROIs were drawn using the ‘interactive thresholding’ and ‘logical expression’ tools along the inside margin of the cell, and separated into individual ROIs using the ‘split by a line’ tool. A minimum of 550 cells were analyzed per sample, for a total of 3,203.

### Calculations

Cells that had low ion count recovery resulting in isotope ratios with a Poisson error greater than 0.1 were excluded from any further analysis. C_net_% and N_net_% were calculated using the isotope fraction (a; at %) of a particular element (X):

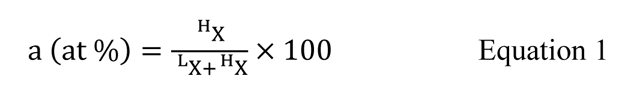

where the superscripts H and L denote the heavier and lighter isotope, respectively.

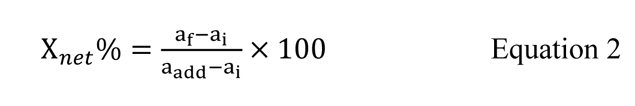

where a_i_ and a_f_ denote the cellular isotope fraction (at %) at the initial and final time point, respectively; and a_add_ is the isotope fraction (at %) of the isotope label (modified from (Popa et al. 2007) as in (Dekas et al. 2019)(Parada et al., 2023)). The isotope fraction of the isotope label at the start of the incubation was calculated using isotope mass balance, the sediment:seawater slurry ratio, the *in-situ* porewater geochemical and porosity data obtained from the IODP LIMS database, assuming an isotope composition of natural abundance for the geochemical species present *in situ* (Rosman and Taylor, 1998), and assuming a porewater bicarbonate concentration of typical seawater (2.06 mM). For the cellular isotope fraction at the initial time point, we used natural abundance, single-cell isotope data from hydrothermally-influenced benthic microbial cells from the East Pacific Rise (Meyer, unpublished data). A cell was identified as active in ammonium or bicarbonate assimilation if it had an N_net_% or C_net_% value that was greater than the mean plus three times the standard deviation of the value for these natural abundance cells (as in (Dekas et al Frontiers 2019)): 0.08% for N_net_ and 0.32% for C_net_. The Gini Coefficient was calculated as in (Arandia-Gorostidi et al., 2023). A two sample, Mann-Whitney (MW) test with a Bonferroni correction for multiple hypotheses was used to test statistical significance, unless otherwise specified. The significance of a linear correlation was tested using the Pearson correlation coefficient (*r*) with a Bonferroni correction. All data (including negative values) were included in the calculations.

## Results and Discussion

### Lithology, sedimentation rate, and geothermal gradient

The sediments at U1550B were mainly biogenic, laminated olive-gray diatom clays or siliciclastic fine (clay) to coarse-grained (sand and silt) components either mixed in homogenous layers or segregated into depositional layers. At the bottom of Hole U1550B, at ∼170 mbsf, a hypabyssal igneous lithology was recovered with a largely doleritic texture. The occurrence of the calcareous nannofossil *Emiliania huxleyi* from the top to the bottom of the hole dates the sediment sequence to (Holocene–)late–middle Pleistocene, or younger than 0.29 Ma, with an estimated average sedimentation rate of >692 m/My (>69.2 cm/ky). Nine *in-situ* formation temperature measurements indicate a linear geothermal gradient of 225°C/km (Teske et al. 2021a) (Figure 2).

**Figure 2:**
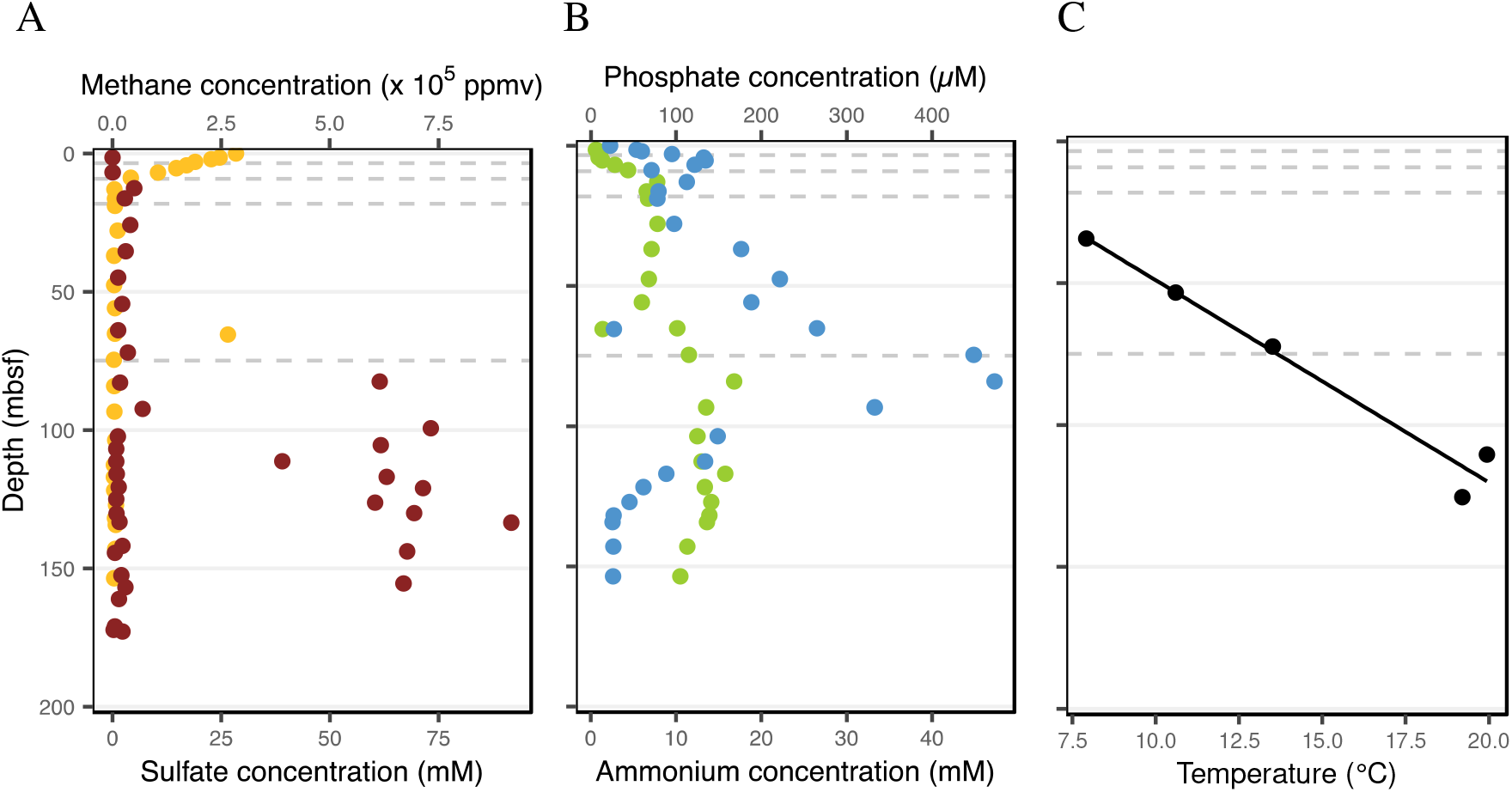
*In situ* geochemical and thermal data at U1550B. A) Porewater concentrations of methane (brown), sulfate (yellow), B) ammonium (green) and phosphate (blue) relative to sediment depth. C) Downhole temperature measured *in situ*. Gray shaded area indicates the 95% confidence interval of the regression line. All data obtained from the IODP LIMS database.

### Solid and porewater geochemistry

At U1550B, the mean organic carbon content was 2.0 ± 0.9 wt% (± 1SD) and mean inorganic carbon content was 0.9 ± 0.5 wt%. The sulfate methane transition zone occurred around 40–50 mbsf, where the methane concentrations sharply increased and porewater sulfate concentrations decreased to the limit of detection. From 0–50 mbsf, porewater ammonium and phosphate concentrations increased with depth (Figure 2). C_2_–C_6_ hydrocarbons were only detectable >100 mbsf (Teske et al. 2021a).

### Proportion of active cells and physicochemical controls on microbial anabolic activity in the subsurface

With depth into the subsurface, the availability of energy and most nutrients decreases (Jørgensen and Boetius 2007). Hence, we expected single-cell microbial anabolic activity to decrease with depth. Indeed, we observe that mean total anabolic activity – calculated as the average single-cell net ^15^N-ammonium assimilation (N_net_%) – decreased with depth (Table 1, Figure 3). But median N_net_% were relatively consistent for samples between 9.1 and 74.9 mbsf (2H-3, 3H-3, and 9H-3) at 0.1–0.2%. At these deeper depths, the low single-cell ammonium uptake suggests that the cells are mostly maintaining their biomass, instead of actively dividing. If a microbial community is doubling through binary fission during the incubation, daughter cells should have N_net_% equal to or greater than 50%. However, we only see cells with N_net_% greater than 50% in the two shallowest samples (≤9.14 mbsf; 1H-3 and 2H-3), suggesting that successful doubling only occurred in the shallowest cores, even after ∼10 months. This is supported by cell count data which shows cell numbers did not increase over the course of the incubation in these deeper incubations, and only slightly in the most shallow sample (Table S1).

**Figure 3:**
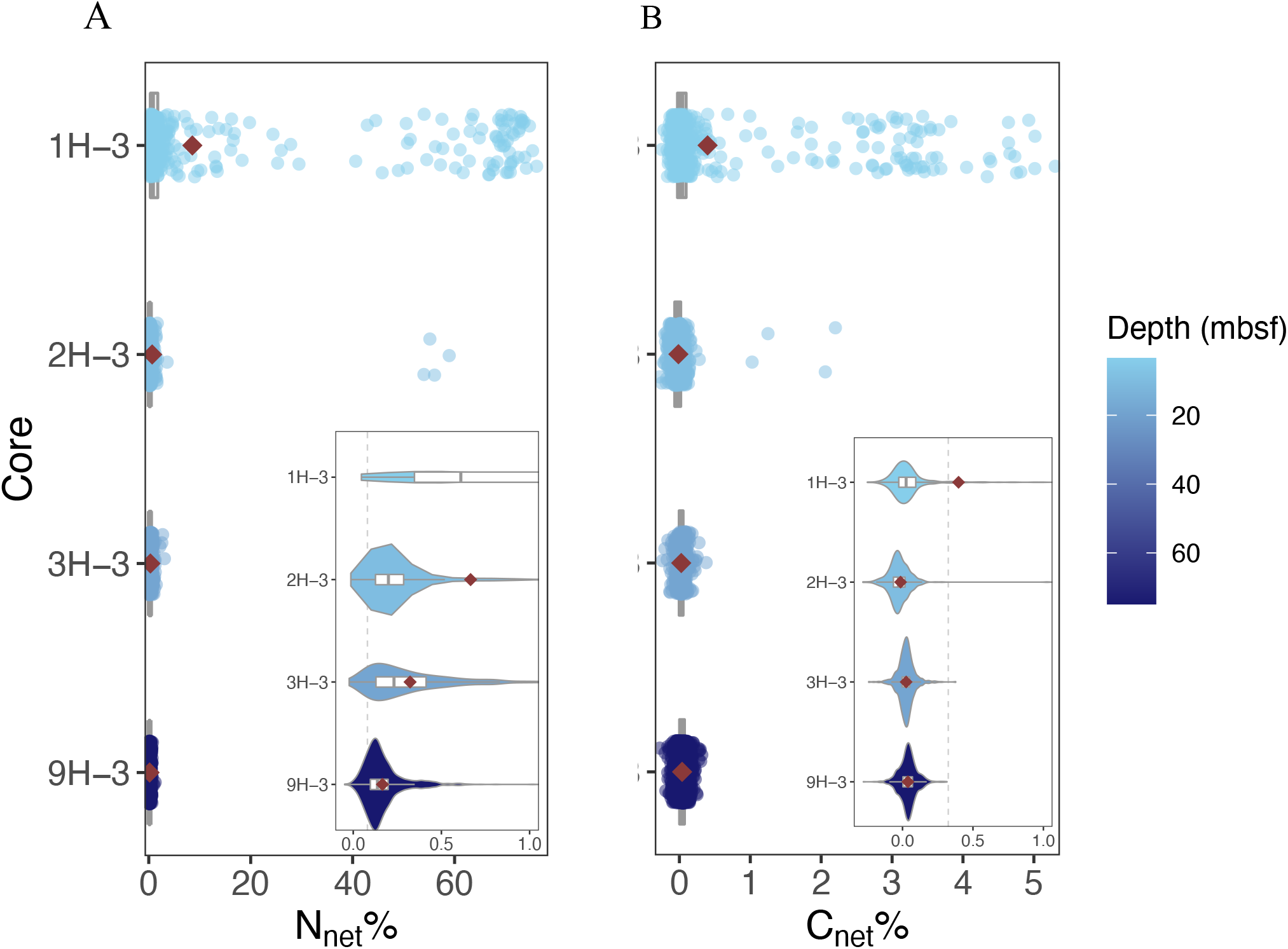
Distribution of A) N_net_% and B) C_net_% values with sediment depth at U1550B for incubations amended with ^15^N-ammonium and ^13^C-bicarbonate. Marker = individual cells. Red diamond = mean. Box and whisker diagram shows the median and quartiles. Violin plots show the probability density. Insets show same data on an adjusted scale, with the threshold for significant enrichment shown with the dotted gray line.

Although single-cell activity rates were low, we nonetheless find that most cells are active, with 100% active in the shallowest sample (3.4 mbsf; 1H-3) and still 83% in the deepest sample (74.9 mbsf; 9H-3) (Table 1, Figure 5). These percentages are high compared to other deep subsurface samples also incubated with ^15^N-ammonium and ^13^C-bicarbonate, where active cells were a minority, albeit after shorter incubation periods: 22% in 219-m-deep sediments from the northwestern Pacific after 65 days (Morono et al. 2011) and 38% in 75-m-deep sediments from the oligotrophic South Pacific Gyre after 21 days (Morono et al. 2020). But our values are comparable to shallower horizons in the South Pacific Gyre (82–94% for cores between 2–69 mbsf) after 21 days and longer incubations of the deeper samples (100% of cells at 75 mbsf after 557 days) (Morono et al. 2020).

We determined the significance of linear correlations between N_net_% and the measured physicochemical variables (Table 2). We found that Pearson correlation coefficients were significant (p ≤ 0.05) for all tested variables except porewater potassium ion concentrations. The geothermal gradient at U1550B is an order of magnitude steeper than the average geothermal gradient (225°C/km (Teske et al., 2021) versus 15–30°C/km within the upper 100 km (Earle 2015) respectively; Figure 2). Microbial activity generally increases with temperature until proteins begin to denature (Sorokin 1960; Arnosti et al. 1998; Kallmeyer and Boetius 2004), but at Site U1550, we do not observe a positive relationship between temperature and N_net_% despite being well below this threshold (Somero 1995). In fact, we observe a small but significant negative relationship between temperature and N_net_% and temperature (r = −0.19; p ≤ 0.005), potentially reflecting a higher maintenance energy at high temperature due to accelerated turnover of biomolecules. We see that N_net_% is most strongly correlated with sulfate (r = 0.33; p ≤ 0.005), the most important electron acceptor in the marine benthic environment due to its abundance (Bradley et al. 2020). Thus, energy may be the most important control on activity at this site.

**Table 2:** Table showing the Pearson correlation coefficients and p-values of single-cell N_net_% and *in situ* porewater geochemistry, temperature, and porosity at U1550B. All incubations were amended with ^15^N-ammonium and ^13^C-bicarbonate. p-values have been adjusted for multiple hypothesis testing using a Bonferroni correction.

| Variable | Correlation coefficient | p-value |
| --- | --- | --- |
| Sulfate | 0.33 | <0.005 |
| Calcium | 0.30 | <0.005 |
| Bromide | -0.28 | <0.005 |
| Lithium | -0.25 | <0.005 |
| Porosity | 0.23 | <0.005 |
| Methane | -0.23 | <0.005 |
| Propane | -0.21 | <0.005 |
| Chloride | -0.20 | <0.005 |
| Temperature | -0.19 | <0.005 |
| Ethene | -0.19 | <0.005 |
| Magnesium | -0.18 | <0.005 |
| Ethane | -0.18 | <0.005 |
| Propene | -0.16 | <0.005 |
| Phosphate | -0.15 | <0.005 |
| Sodium | 0.09 | <0.005 |
| Strontium | -0.06 | 0.014 |
| Potassium | 0.01 | 1.000 |

### Intracommunity heterogeneity in single-cell anabolic activity

Cells in both isogenic, laboratory cultures (Kopf et al. 2015; Schreiber et al. 2016; Calabrese et al. 2019, 2021) and mixed community, environmental samples (Vasquez-Cardenas et al. 2015; Arandia-Gorostidi et al. 2017; Dekas et al. 2019) can show a large range in single-cell activity. In pure cultures, the heterogeneity in activity might be a result of stochastic gene expression, cell growth stage, cell-to-cell interactions, and the co-occurrence of genotypes and ecotypes (Musat et al. 2008; Woebken et al. 2012; Berry et al. 2013; Berthelot et al. 2019; Calabrese et al. 2021). In mixed cultures and natural samples, activity heterogeneity is also affected by genetic heterogeneity. Here we explored the relationship between intracommunity heterogeneity in activity and sediment depth by calculating the Gini coefficient (Gini 1921) using our cell-specific ^15^N-ammonium uptake rates as in (Arandia-Gorostidi et al., 2023). The Gini coefficient ranges from zero to one, with higher values representing more heterogeneity in distribution, i.e., unevenness.

We find that the distribution of single-cell activity is increasingly even with increasing subsurface depth (Figure 4). This differs from previous studies that have shown that both nutrient limitation and slower growth rates tend to increase heterogeneity in microbial activity in pure cultures (Kopf et al. 2015; Zimmermann et al. 2015, 2018; Schreiber et al. 2016). However, in environmental samples like at U1550B, many potential drivers of activity heterogeneity change simultaneously, and thus trends in single-cell activity distribution may not be as simple and predictable as those in laboratory, isogenic populations. The decreasing heterogeneity in activity with depth observed here might be a product of decreasing genetic heterogeneity; anoxic marine sediments generally decrease in taxonomic richness with subsurface depth globally (Hoshino et al. 2020). Additionally, in some laboratory bacterial cultures, single-cell activity decreases in heterogeneity over time (Calabrese et al. 2021), potentially due to metabolism optimization (Goel et al. 2012). Sediment age increases with depth into the subsurface, and thus, metabolism optimization over time might explain the decrease in activity heterogeneity with depth and sediment age (Figure 4).

**Figure 4.**
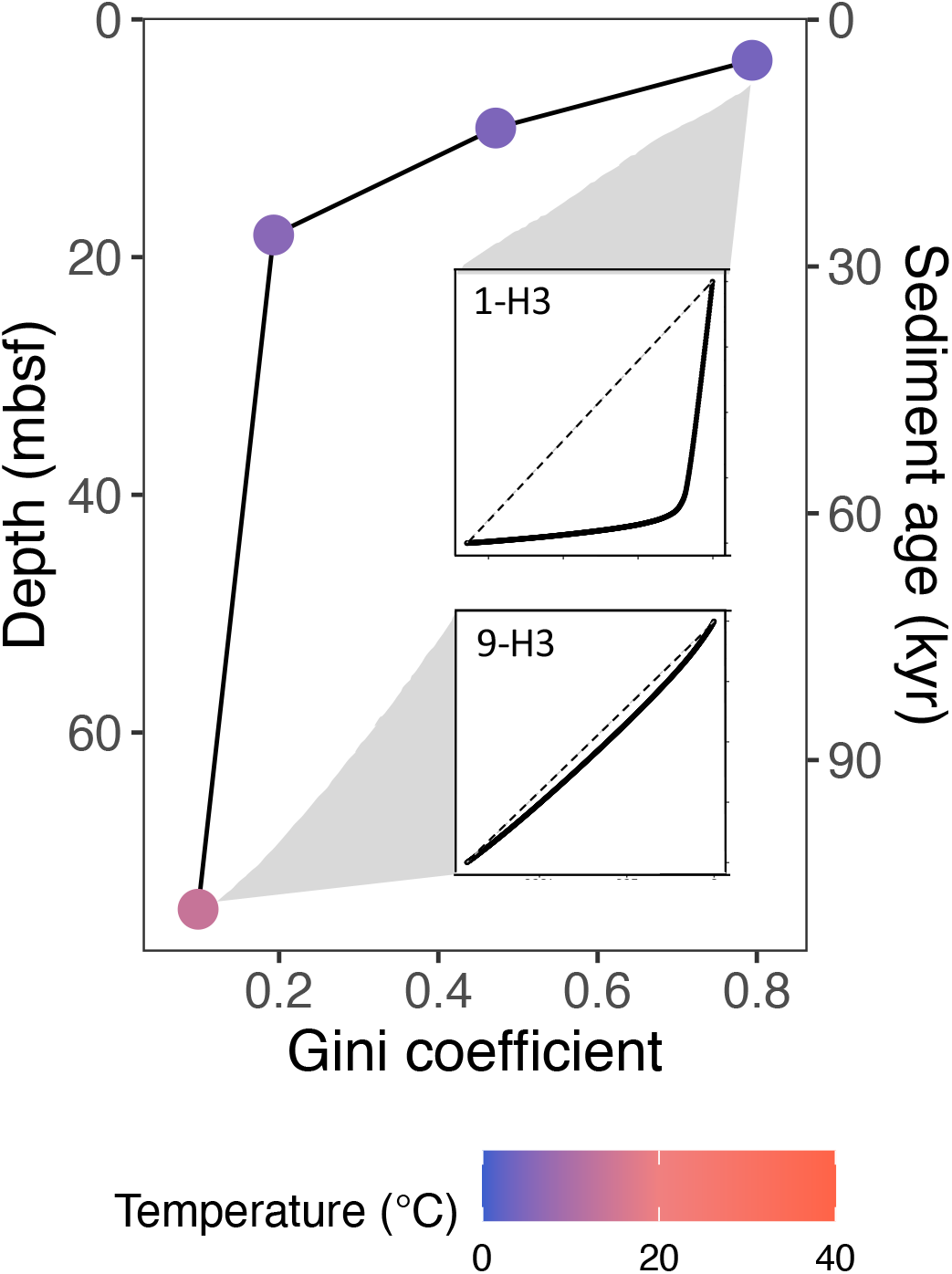
The relationship between phenotypic heterogeneity and sediment depth and age. Single-cell nitrogen isotope compositions were used to calculate Gini coefficients, a measure of inequality, where higher values = greater unevenness. Insets show Lorenz curves for the shallowest and deepest samples; for these, cumulative share of total summed N_net_% is represented on the y-axis and cumulative share of cells from least to most active is represented on the x-axis. The dashed 1:1 line indicates a community with equal activity across its members. All incubations were amended with ^15^N-ammonium and ^13^C-bicarbonate.

### Characterizing carbon uptake metabolisms

The deep subsurface might be dominated by heterotrophs as indicated by the widespread occurrence of genes associated with organic matter degradation (Biddle et al. 2006; Lloyd et al. 2013; Orsi et al. 2013; Baker et al. 2015; Marshall et al. 2018; Orsi 2018; Zinke et al. 2019; Wasmund et al. 2021). But marine benthic hydrogenotrophic methanogens (Beulig et al. 2018) and some methanotrophs (Kellermann et al. 2012) are known to assimilate inorganic carbon and genetic evidence suggests they are widespread in the subsurface (Orcutt et al. 2011). However, the presence of genes does not equate activity. Direct uptake of inorganic carbon has been observed in some subsurface samples, including in the South Pacific Gyre (Morono, 2020), however, autotrophy and inorganic carbon fixation by heterotrophs were not distinguished. A quantitative estimate of heterotrophic to autotrophic cells is therefore unknown in the deep subsurface. We wanted to determine this in the deep-subsurface at Guaymas Basin, hypothesizing that it was dominated by heterotrophs due to its high organic carbon content (mean = 2.0 wt% at Hole U1550B) and the potential of pyrolysis products advecting from below. However, diverse methanogens have been detected by PCR-based surveys at Guaymas Site 1550 (Hinkle, preprint), suggesting the genetic potential for autotrophy.

We can use the single-cell uptake of ^13^C-bicarbonate relative to ^15^N-ammonium as an indicator of the primary carbon uptake metabolism, i.e. to differentiate between autotrophs and heterotrophs (Dekas et al., 2019). This approach identifies heterotrophs by a process of elimination: cells active in ^15^N-ammonium uptake with no (or low) ^13^C-bicarbonate uptake must be deriving the bulk of their carbon from organic sources, similar to the concept of “lipid dual SIP” (Kellermann et al., 2012; Wegener et al., 2012). Although not a direct measure of organic carbon uptake, it is likely an inclusive indicator of heterotrophy because it is not dependent on a heterotroph’s ability to assimilate a particular organic substrate. In this case, since no organic compounds were added, heterotrophy would be supported by the existing pool of sedimentary organic carbon, as observed previously to occur in subsurface samples (Morono et al., 2020). Using previously established thresholds (Dekas et al. 2019), a cell is operationally categorized as an autotroph if it obtains more than 50% of its carbon from bicarbonate (2×C_net_% > N_net_%) while a heterotroph obtains more than 50% of its carbon from organic sources (N_net_ % > 2×C_net_%). An organism that is active (N_net_% > the limit of detection) but does not assimilate any ^13^C-bicarbonate (C_net_% < the limit of detection) obtains 100% of its carbon from organic sources (Figure 5). Thus, plotting C_net_% and N_net_% together can help classify the dominant carbon uptake metabolism.

**Figure 5:**
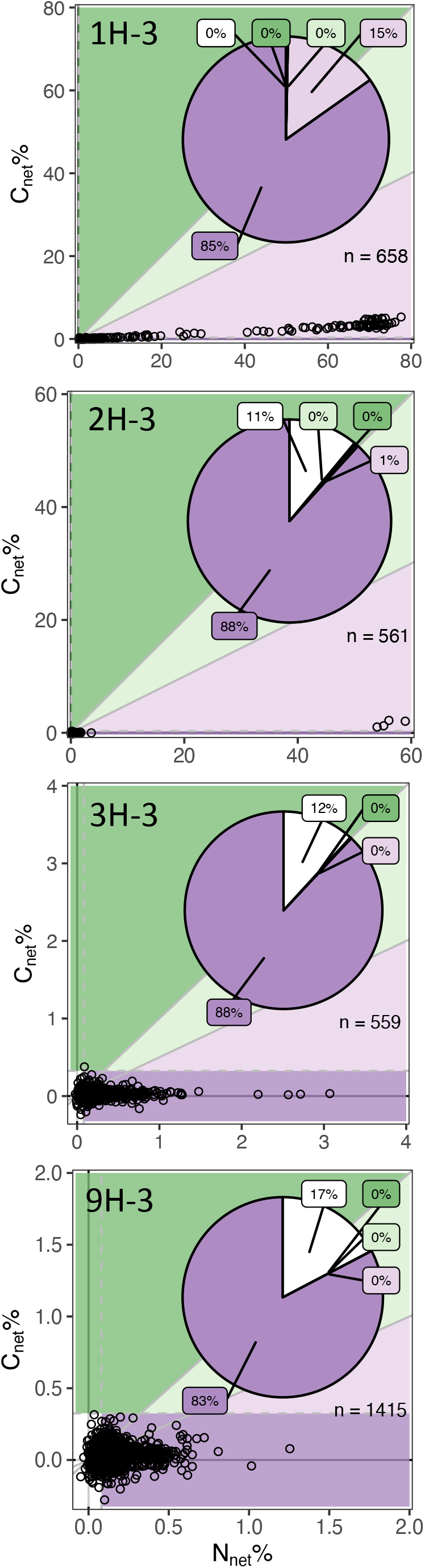
Metabolic characterization of individual cells from each U1550B core based on relative assimilation of ^13^C-bicarbonate and ^15^N-ammonium. Markers = individual cells. Colors indicate metabolic regimes as defined in (Dekas *et al*., 2019); white: below the limit of detection (inactive cells); purple: >50% of new C from organic sources (heterotrophy); green: >50% of new C from bicarbonate (autotrophy). Dark purple and green regions indicate 100% of new carbon from organic and inorganic sources, respectively. Dashed grey lines indicate the limit of detection, i.e., 3X the standard deviation from the mean of isotope unlabeled cells. Note the axes are variable to optimize visualization. Inset pie charts show the proportion of inactive cells, heterotrophs, and autotrophs in the same color scheme.

Most active cells at U1550 did not assimilate a detectable amount of ^13^C-bicarbonate, indicating widespread heterotrophy in these samples. However, in our two shallowest samples (3.4 mbsf [1H-3] and 9.1 mbsf [2H3]), we did find that 15% and 1 % of cells, respectively, have a C_net_% up to 5% (Figure 3). Normalizing the C_net_% value to the N_net_% value per cell indicates that this assimilation of inorganic carbon comprises a small percentage of the cell’s total carbon intake, and that these cells are therefore heterotrophs (Figure 5). Interestingly, we did not detect any ^13^C-bicarbonate uptake within the sulfate methane transition zone (74.9 mbsf; 9H-3), where we might expect methanotrophs. Together, for all depths at the Guaymas Basin northern axial trough, heterotrophs comprised 100% of the active microbial community.

### Quantifying the magnitude of inorganic carbon fixation by heterotrophs

All heterotrophs – from microorganisms to humans – fix some inorganic carbon (Braun et al. 2021). Heterotrophs have carboxylating enzymes that can synthesize fatty acids, amino acids, vitamins, and nucleotides, as well as perform leucine assimilation and anaplerosis (Evans and Slotin 1940; Krebs 1941; Kornberg and Krebs 1957; Kornberg 1965; Hartman et al. 1972; Schink 2009). In these cases, inorganic carbon is a “co-substrate” that is used to extend an existing organic substrate by a single carbon unit (Erb 2011). Anaplerosis is the most quantitatively important inorganic carbon fixation pathway in all organisms, and it replenishes intermediates in the tricarboxylic acid cycle (TCA). Products of the TCA cycle are used as components of essential macromolecules like amino acids (Krebs 1941).

The magnitude of heterotrophic inorganic carbon fixation is still debated. Most data originate from isogenic, laboratory cultures using carbon isotope labeling, where heterotrophs obtain between 1 and 13% of their carbon from CO_2_ or bicarbonate (Perez and Matin 1982; Roslev et al. 2004; Hesselsoe et al. 2005; Feisthauer et al. 2008; Spona-Friedl et al. 2020). In the environment, anaplerotic reaction rates are more difficult to quantify because heterotrophs and autotroph usually co-exist. Thus, previous environmental studies have tried to measure bulk, inorganic carbon uptake rates where there is no molecular evidence of chemoautotrophs and their related genes, or where the oxic environment lacks potential inorganic electron donors (e.g. hydrogen sulfide, ammonium, hydrogen, etc.). For example, in soils presumed to be dominated by the activity of heterotrophs, dark inorganic carbon fixation rates were 0.04–39% of the overall respiration rate or microbial production (Miltner et al. 2004, 2005; Santruckova et al. 2005; Šantrůčková et al. 2018; Akinyede et al. 2020; Spohn et al. 2020). But the large range in estimates might still be due to accidental inclusion of chemoautotrophy in the rate of inorganic carbon fixation, not only heterotrophic inorganic carbon fixation. Thus, there is a real need for an improved, quantitative approach to measure inorganic carbon fixation by heterotrophs in mixed microbial communities in the environment.

Here, our dual isotope tracer and single-cell approach allows us to quantify the portion of cellular carbon derived from inorganic sources for each cell, and therefore the contribution of inorganic carbon to heterotrophic biomass. We observe that C_net_% and N_net_ % follow a linear regression across all subsurface depths (R^2^ = 0.94, p ≤ 0.001; Figure 6). The slope of the line indicates that on average, the heterotrophs at U1550B obtained 5% of their carbon from inorganic sources. Sample preparation for nanoSIMS causes a slightly greater decrease in ^13^C enrichment than ^15^N enrichment (specifically cellular fixation), and thus our percentage is a lower threshold (Musat et al. 2014; Woebken et al. 2015; Meyer et al. 2021). Notably, although this community turned out to be entirely comprised of heterotrophs, the selectivity of this method would warrant it equally effective in communities of mixed heterotrophs and autotrophs. In that case, a regression could be generated of the heterotrophic cells alone to determine their rate of inorganic carbon fixation. As an example, and for comparison, we here applied this approach to previously published nanoSIMS data from cells specifically identified as heterotrophs in the pelagic Pacific Ocean (150 m water depth at the San Pedro Ocean Times Series, SPOT) (Dekas et al Frontiers 2019). We detected inorganic carbon fixation by heterotrophs in this metabolically mixed pelagic community, as well, but less than observed in the subsurface heterotrophs (1% of their total carbon versus 5%; Figure 6). Although more observations are needed to draw any comparative conclusions, this suggests that heterotrophic inorganic carbon assimilation may be more significant in subsurface sediments than other marine habitats.

**Figure 6:**
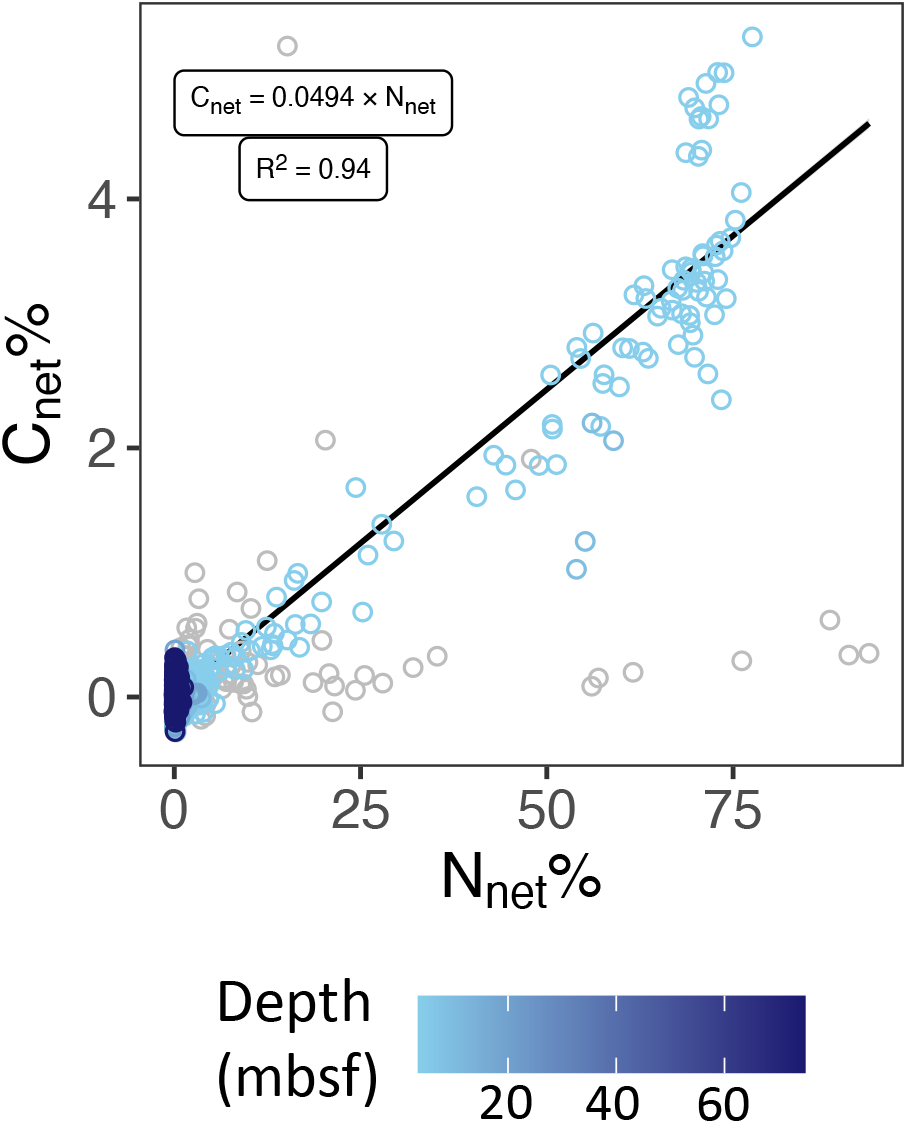
Net N and C uptake (N_net_% and C_net_%) for cells incubated with ^15^N-ammonium and ^13^C-bicarbonate for all cells at U1550B (cores 1H-3, 2H-3, 3H-3, and 9H-3 together) (blue circles). For comparison, heterotrophic sea surface cells from the Northeast Pacific Ocean are included as gray circles (Dekas et al, Frontiers 2019). Each circle displays data from an individual cell. The equation of the line and R^2^ value for the U1550B cells are indicated on the plot, those for the pelagic data are as follows: C_net_ = 0.0095 x N_net_, R^2^ = 0.12.

We believe our estimates are the first direct measurements of inorganic carbon fixation within a natural population of individually identified heterotrophic cells within a complex environmental sample. Our estimates fall within the range of those found previously in laboratory cultures (Perez and Matin 1982; Roslev et al. 2004; Hesselsoe et al. 2005; Feisthauer et al. 2008; Spona-Friedl et al. 2020) and environmental samples using a bulk approach (Miltner et al. 2004, 2005; Santruckova et al. 2005; Šantrůčková et al. 2018; Akinyede et al. 2020; Spohn et al. 2020). This validation supports previously proposed upper estimates that 2–4 Pg C of Earth’s total living biomass is from heterotrophic inorganic carbon fixation (Braun et al. 2021). It also highlights the specific role of heterotrophic inorganic carbon fixation in the subsurface.

## Conclusion

The hydrothermal deep subsurface at the Guaymas Basin axis contains active cells that are contributing to the biogeochemical cycling of nitrogen and carbon. Their activity decreases with depth, following the availability of electron acceptors like sulfate, and do not positively correlate with temperature. Single-cell activity is more evenly distributed between community members with subsurface depth, potentially due to metabolic optimization over time. Heterotrophs dominate, and they obtain on average 5% of their carbon from inorganic sources. The deep biosphere in the Guaymas Basin plays an integral role in subsurface carbon cycling, by assimilating organic carbon and reducing the benthic flux of carbon dioxide through heterotrophic inorganic carbon fixation.

## Acknowledgments

We would like to thank the shipboard crew, operational team members, and shipboard scientists of IODP expedition 385. In particularly, we would like to thank the microbiology team on IODP expedition 385 for their insightful discussions. We thank members of the Dekas laboratory for their helpful feedback and comments. We also thank Megumi Becchaku and Takeshi Terada for technical assistance. This study was supported by a US Science Support Party (USSSP) post-expedition activity award (102C(GG009393-04)) to N.M., a USSSP Schlanger fellowship to N.M., a National Science Foundation CAREER award to A.D. (2143035) and JSPS KAKENHI Grants JP19H00730, JP22K18426, JP23H00154 to YM.

---

**Table S1.** Cell densities before and after incubation.

